## Supplementary Material for "HIV-1 promoter is gradually silenced when integrated into *BACH2*"

**Supplementary Table 1:** Primers used to amplify the *BACH2* and AAVS1 homologous arms, to amplify and sequence the junction between the vector LTatCL[M] and *BACH2* or AAVS1, to test for mono- or bi-allelic integration, and to amplify the vector LTatCL[M].

| **Primer name** | **Primer sequence (5‘-3‘)** | **Description** |
|---|---|---|
| BsrGI_BACH2_1_5fw | NNNN*TGTACA*CACTTTCGTGGAGCTG TTGTTAG | BsrGI overhang, forward primer, cloning of *BACH2*_i5 5` homologous arm |
| PacI_BACH2_1_5rc | NNNN*TTAATTAA*TTCTGGGTTCTGTTA CTCAC | PacI overhang, reverse primer, cloning of *BACH2*_i5 5` homologous arm |
| AscI_BACH2_1_3fw | NNNN*GCGCGCC*TAGCATGGAGGAGT ATGACAG | AscI overhang, forward , cloning of *BACH2*_i5 3` homologous arm |
| SpeI_BACH2_1_3rc | NNNNNN*ACTAGT*GCATTAGAAAGTGT TTCCAATCC | SpeI overhang, reverse primer, cloning of *BACH2*_i5 3` homologous arm |
| BsrGI_BACH2_2_5fw | NNNN*TGTACA*GTAACAGTGCTCTCTA TACATTATCC | BsrGI overhang, forward primer, cloning of *BACH2*_i2 5` homologous arm |
| PacI_BACH2_2_5rc | NNNNNN*TTAATTAA*CTATGGTTTCACC TTTTCTGGATATTTC | PacI overhang, reverse primer, cloning of *BACH2*_i2 5` homologous arm |
| AscI_BACH2_2_3fw | NNNN*GGCGCGCC*ACGCAGGAAGCAG AGAAGTG | AscI overhang, forward primer, cloning of *BACH2*_i2 3` homologous arm |
| SpeI_BACH2_2_3rc | NNNNNN*ACTAGT*CTACTTTGATATTT GAAGGATAGATTCC | SpeI overhang, reverse primer, cloning of *BACH2*_i2 3` homologous arm |
| BsrG1_AAVS1_5arm_f w | NNNN*TGTACA*GGTCCTGCTTTCTCTG ACCTGC | BsrGI overhang, forward primer, cloning of AAVS1 5` homologous arm |
| Pac1_AAVS1_5arm_rc | NNNN*TTAATTAA*TGTCCCTAGTGGCCC CACTGTG | PacI overhang, reverse primer, cloning of AAVS1 5` homologous arm |
| Asc1_AAVS1_3arm_fw | NNNN*GGCGCGCC*GGATTGGTGACAG AAAAGCCCCATC | AscI overhang, forward primer, cloning of AAVS1 3` homologous arm |
| Spe1_AAVS1_3arm_rc | NNNNNN*ACTAGT*GTCTGAAGAGCAG AGCCAGGAACC | SpeI overhang, reverse primer, cloning of AAVS1 3` homologous arm |
| geneB21_fw | ATACAAGGAAACCACAGCCTTCTGG | outer/inner PCR of junction, mono- or bi-allelic integration, and vector amplification and sequencing of LTatCL[M]/*BACH2*_i5s and LTatCL[M]/*BACH2*_i5c |
| REV_Tat_rc | TCTAGTCTAGGATCTACTGGCTCC | outer PCR of junction LTatCL[M]/*BACH2*_i5c, and inner PCR of junction LTatCL[M]/*BACH2*_i2c |
| nB21_fw | CAGCAACATAAGCATCCCAAGTTGA TG | inner PCR of junction LTatCL[M]/*BACH2*_i5c, and inner PCR of junction LTatCL[M]/*BACH2*_i5s |
| 5’LTRIII | TGTGGTAGATCCACAGATCAAG | outer PCR of junction LTatCL[M]/AAVS1_c, and inner PCR of junction LTatCL[M]/*BACH2*_i5c and LTatCL[M]/AAVS1_c |
| KYL-INS3LTR-1897fw | GTCAACATCAAGTTGGACATCACC | outer PCR of junction LTatCL[M]/AAVS1_s, and inner PCR |

| | | |
|---|---|---|
| | | of junction LTatCL[M]/*BACH2*_i5s |
| PolyA_RO_rc | TGTGTCTAGAGCTCGAGCATGC | inner PCR of junction, and vector amplification and sequencing of LTatCL[M]/*BACH2*_i5s, LTatCL[M]/*BACH2*_i2s and LTatCL[M]/AAVS1_s |
| geneB22_fw | ATACAAGGAAACCACAGCCTTCTGG | outer PCR of junction, mono- or bi-allelic integration, vector amplification and sequencing of LTatCL[M]/*BACH2*_i2s and LTatCL[M]/*BACH2*_i2c |
| SigmaP1_fw | GGCCCTGGCCATTGTCACTT | outer PCR of junction LTatCL[M]/AAVS1_s and LTatCL[M]/AAVS1_c |
| LJ_AAVS1_fw3 | CTTTGAGCTCTACTGGCTTCTGC | inner PCR of junction and mono- or bi-allelic integration LTatCL[M]/AAVS1_s and LTatCL[M]/AAVS1_c |
| A.I_BACH2.1_3'arm_rc | AGACTGGCTGTCATACTCCTCCATG | mono- or bi-allelic integration of LTatCL[M]/*BACH2*_i5s and LTatCL[M]/*BACH2*_i5c |
| AI_AAVS1_3'arm1Rv | GCCTAAGGATGGGGCTTTTCTG | mono- or bi-allelic integration of LTatCL[M]/AAVS1_s and LTatCL[M]/AAVS1_c |
| A.I_BACH2.2_3'arm_rc | GAAACCACTTCTCTGCTTCCTGC | mono- or bi-allelic integration of LTatCL[M]/*BACH2*_i2s and LTatCL[M]/*BACH2*_i2c |
| KYL_5LTRFL_1121rc | GGCACGCGTCTAATCGAATGG | vector amplification and sequencing of LTatCL[M]/*BACH2*_i5s, LTatCL[M]/*BACH2*_i5c, LTatCL[M]/*BACH2*_i2s and LTatCL[M]/*BACH2*_i2c |
| PolyA_fw | GCATGCTCGAGCTCTAGACACA | vector amplification and sequencing of LTatCL[M]/*BACH2*_i5s, LTatCL[M]/*BACH2*_i5c, LTatCL[M]/*BACH2*_i2s and LTatCL[M]/*BACH2*_i2c |
| Rev_Tat_fw | GGAGCCAGTAGATCCTAGACTAGA | vector amplification and sequencing of LTatCL[M]/*BACH2*_i5s, LTatCL[M]/*BACH2*_i5c, LTatCL[M]/*BACH2*_i2s and LTatCL[M]/*BACH2*_i2c |
| gBACH2_1_Fw | CACCGATACTCCTCCATGCTATTCT | *BACH2*_i5 gRNA (cloning of guide sequence into pX458) |
| gBACH2_1_rc | AAACAGAATAGCATGGAGGAGTATC | gRNA of *BACH2*_i5 (cloning of guide sequence into pX458) |
| gBACH2_2_fw | CACCGAAAGGTGAAACCATAGACGC | gRNA of *BACH2*_i2 (cloning of guide sequence into pX458) |
| gBACH2_2_rc | AAACGCGTCTATGGTTTCACCTTTC | gRNA of *BACH2*_i2 (cloning of guide sequence into pX458) |
| gAAVS1_fw | CACCGGGGCCACTAGGGACAGGAT | gRNA of AAVS1 (cloning of guide sequence into pX458) |
| gAAVS1_rc | AAACATCCTGTCCCTAGTGGCCCC | gRNA of AAVS1 (cloning of guide sequence into pX458) |

| | | |
|---|---|---|
| BaEx7-Fw | GAACCAACTCCAGTGACGAATCC | mRNA quantification downstream of *BACH2*_i2 and *BACH2*_i5 |
| BaEx8-Rv | CTAACTGTTCTGAGGTTAGCTTGTGC | mRNA quantification downstream of *BACH2*_i2 and *BACH2*_i5 |
| Mf45 | TCGACAGTSAGCCGCATCTT | mRNA quantification of Glycerinaldehyd-3-phosphat-Dehydrogenase (GAPDH) |
| Mf46 | GGCAACAATATCCAGTTTACCAG | mRNA quantification of Glycerinaldehyd-3-phosphat-Dehydrogenase (GAPDH) |

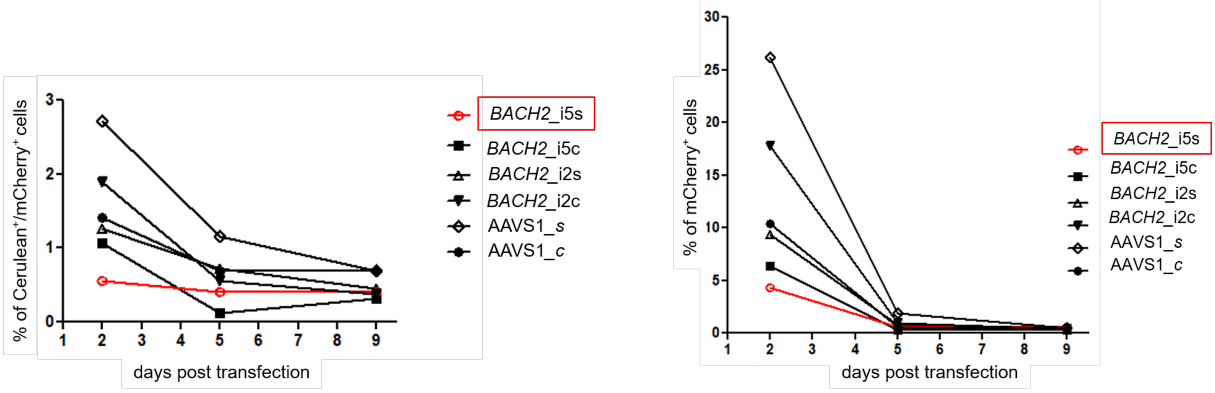


**Supplementary Figure 1: FACS analysis of Cerulean$^+$/mCherry$^+$ and single mCherry$^+$ expression 2, 5, and 9 days post transfection.** Percentage of Cerulean$^+$/mCherry$^+$ (left panel) and single mCherry$^+$ (right panel) expressing cells over time in all six cell variants each, transfected with one of the six vectors LTatCL[M], are shown for one exemplary experiment. Each symbol represents one cell variant, open symbols represent cells in which LTatCL[M] is integrated in the same transcriptional orientation of the gene, closed symbols represent cells in which LTatCL[M] is integrated in the convergent transcriptional orientation of the gene. The experiment was carried out three times independently. The *in vivo* observed preferential HIV-1 integration loci in *BACH2, BACH2*_i5s, is highlighted by red boxes.

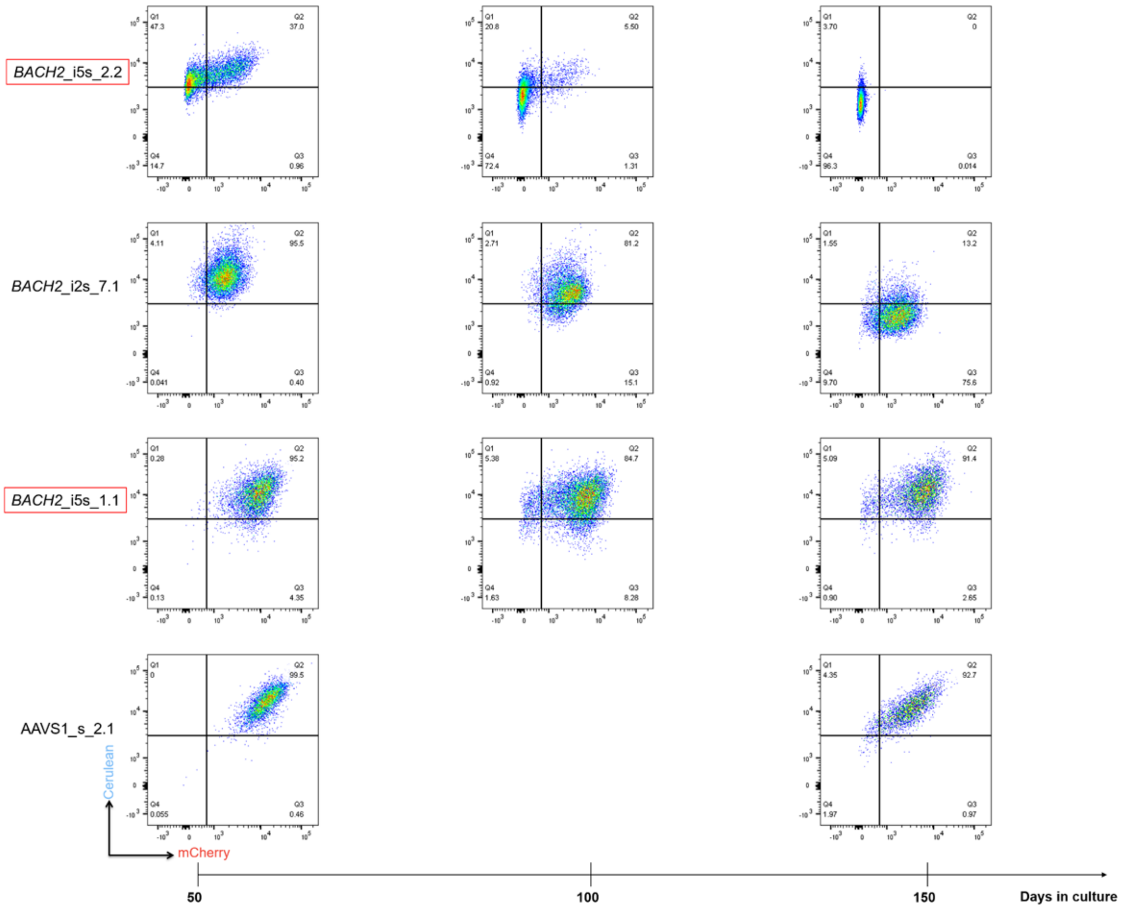


**Supplementary Figure 2: Longitudinal FACS analysis of Cerulean$^{+}$/mCherry$^{+}$ monoclonal cell lines for up to 162 days.** Three time points (50, 100, and 150 days in culture) are depicted for exemplary Cerulean$^{+}$/mCherry$^{+}$ monoclonal cell lines showing different phenotypic changes over time; Cerulean$^{+}$/mCherry$^{+}$ to a Cerulean$^{-}$/mCherry$^{-}$ expressing phenotype depicted for *BACH2*_i5s_2.2, Cerulean$^{+}$/mCherry$^{+}$ to single mCherry$^{+}$ expressing phenotpye depicted for *BACH2*_i2s_7.1 and Cerulean$^{+}$/mCherry$^{+}$ monoclonal cell lines maintaining Cerulean$^{+}$/mCherry$^{+}$ expressing phenotype depicted for *BACH2*_i5s_1.1 and AAVS1_s_2.1. The experiment was carried out two times independently. The *in vivo* observed preferential HIV-1 integration loci in *BACH2, BACH2*_i5s, is highlighted by red boxes.
